# CONET: Copy number event tree model of evolutionary tumor history for single-cell data

**DOI:** 10.1101/2021.04.23.441204

**Authors:** Magda Markowska, Tomasz Cąkała, Błażej Miasojedow, Dilafruz Juraeva, Johanna Mazur, Edith Ross, Eike Staub, Ewa Szczurek

## Abstract

Copy number alterations constitute important phenomena in tumor evolution. Whole genome single cell sequencing gives insight into copy number profiles of individual cells, but is highly noisy. Here, we propose CONET, a probabilistic model for joint inference of the evolutionary tree on copy number events and copy number calling. CONET employs an efficient MCMC procedure to search the space of possible model structures and parameters and utilizes both per-bin and per-breakpoint data. We introduce a range of model priors and penalties for efficient regularization. CONET achieves excellent performance on simulated data and for 260 cells from xenograft breast cancer sample.

## Background

Elucidating tumor evolutionary history is pivotal to a better understanding of carcinogenesis and helps developing new cancer treatments. Copy number alterations (CNAs) are ubiquitous in the genomes of tumors across all types of cancer [1–3], constitute the most common alteration types associated with tumor hypermutability [4], and play an important role as drivers of tumor evolution. In particular, amplifications can activate so called oncogenes, promoting increased growth and other hallmarks of cancer [5], while deletions can disable tumor suppressors [1, 2]. Recently, the technology of single cell DNA sequencing (scDNA-seq) revolutionizes the analysis of tumor cell populations and gives valuable insights into tumor evolution [6]. Modeling copy number evolution of tumors from this data, however, is challenged by the noise, and the fact that copy number events may overlap, invalidating the often-made finite sites assumption. Thus, the methods addressing this problem are still in their infancy [7].

Tumor phylogeny reconstruction from bulk sequencing data has been approached by a multitude of methods, most of which focus on the evolution of single nucleotide variants (SNVs) [8–23]. Bulk tumor DNA sequencing data jointly measures a mixture of millions of cells coming from different subclones and an unknown quantity of healthy cells. Reconstruction of copy number evolution from this data is notoriously difficult, as only the aggregated signal per region is observed [7]. Dedicated methods addressing this problem are challenged by the mixture of different copy number profiles and their proportions in bulk samples or the difficulty of finding appropriate measures of phylogenetic distance between the CNA samples [12, 24–29].

In contrast to bulk sequencing, the more recent scDNA-seq offers the measurement of the genomic sequence at the level of individual cells. As such, it paves the way to computational identification of genomic alterations such as CNAs and SNVs in single genomes, and their evolutionary relationships [30,31]. The single cell measurements, however, are noisy, with major issues being low coverage depth or low coverage uniformity. This calls for dedicated probabilistic approaches to tumor evolution reconstruction from this data [7]. Several approaches have already been proposed for modeling tumor evolution of SNVs from scDNA-seq data [32–36] or jointly from bulk and scDNA-seq [37–39].

A number of scDNA-seq high-throughput techniques, such as DOP-PCR [40–42] and C-PCR-L [43] achieve relatively even coverage distribution along the whole genome of the sequenced cells [7, 44]. Recently, DLP and DLP+ have been proposed as technologies omitting the pre-amplification step, and thus avoiding the amplification bias [45, 46]. These techniques open up a new avenue to copy number calling in single tumor cells, suggesting that also reconstruction of copy number tree evolution models from such type of data is feasible.

Existing copy number calling approaches, however, suffer from high false positive rates [44,47]. In particular, identifying the locations of the borders between genomic segments with different copy number, which are referred to as *breakpoints*, is largely hindered by the inherent noise in the scDNA-seq data. Joint inference of tumor phylogeny from single cells and copy number calling from even coverage scDNA-seq is expected to result in more accurate copy number calls.

There are, in fact, two types of readouts available for copy number evolution inference and copy number calling from raw scDNA-seq data. The first type of readout is the *per-bin* data: sequencing counts in each genomic bin, which, after appropriate normalization and correction for mappability and GC content, indicate the copy number at that bin. The second is the *per-breakpoint* data, given by the change of the counts at the potential breakpoint loci. On the one hand, at a locus where there is no copy number event start or end, we expect no difference between the counts in the bin preceding the locus and the counts in the bin succeeding the locus. On the other hand, at a breakpoint locus we expect a large difference between the counts, and the larger the absolute value of the change, the more evidence for the presence of that breakpoint in the cell.

Recent approaches analyzing copy number changes in single cells from an evolutionary perspective aim at either improving breakpoint and copy number calling from scDNA-seq based on breakpoint trees [48], or modeling cell lineages based on the similarities of copy number profiles between single cells and identifying events carrying fitness advantage [49], or inference of copy number evolution and copy number calling from the binned read counts [50]. None of these approaches, however, infers copy number event trees or identifies copy number states from both per-bin and per-breakpoint data.

Here, we propose a novel approach for Copy Number Event Tree (CONET) inference and copy number calling (Figure 1). CONET fully exploits the signal in scDNA-seq, as it relies directly on both the per-breakpoint and per-bin data. The model jointly infers the structure of an evolutionary tree on copy number events and copy number profiles of the cells, gaining statistical power in both tasks. The nodes of the evolutionary tree are copy number events, which are allowed to overlap. Results on simulated and real breast cancer data demonstrate the excellent performance of CONET in both tasks. Taken together, the proposed approach is a step towards a better understanding of copy number evolution in cancer.

**Figure 1.**
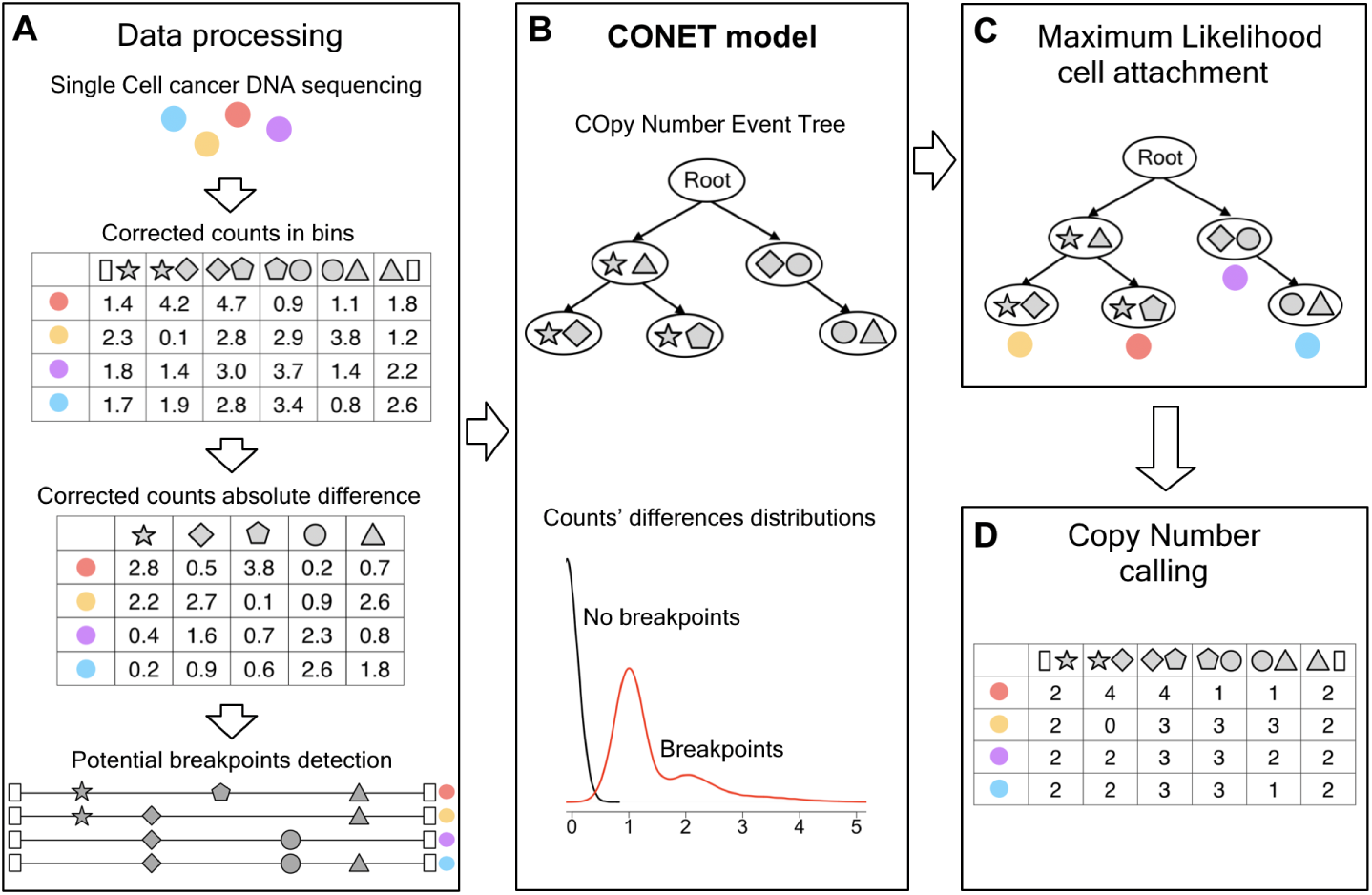
Joint inference procedure of CONET and integer copy numbers. **A** Preparation of sc-DNA-seq data as an input for the model involves calculation of the corrected count absolute differences for all genomic loci (per-breakpoint data) using GC and mappability corrected counts in bins (per-bin data); and defining the set of potential breakpoints. **B** The main CONET inference procedure is an efficient MCMC sampling scheme, searching the space of possible CONET structures and model parameters, using per-breakpoint data for likelihood calculation and per-bin data for model regularisation. **C, D** In the post-processing steps the cells are attached to the final CONET nodes using Maximum Likelihood and the inferred breakpoint history in each single cell, together with per-bin data, are used to determine the cell’s copy number profiles.

CONET implementation is available at https://github.com/tc360950/CONET.

## Results

### Model overview

We propose a joint procedure that efficiently infers the history of copy number events occurring in single cells in the tumor tissue, identifies the presence of breakpoints and estimates the integer copy numbers in regions defined by these breakpoints in each cell from scDNA-seq data (Figure 1). In the first step (Figure 1 A), we process per-bin scDNA-seq data from tumor sample, which needs to be corrected for GC content and mappability, and normalized so that the corrected counts in bins reflect the underlying integer copy numbers with neutral copy number equal to two. To specify the per-breakpoint input data to the CONET model, we calculate the differences between the corrected counts in adjacent bins for each cell, arriving at the corrected count absolute difference matrix (Methods). Finally, we establish the set of candidate breakpoint loci (the definition of this set belongs to the user - see Real data preprocessing).

In the second step (Figure 1 B), we run CONET, a generative probabilistic model for inferring tumor evolution on single cell copy number events. The structure of CONET is defined by a tree, with the root representing a healthy cell with a neutral copy number, and each remaining node corresponding to an evolutionary event interpreted as a copy number change that introduces the occurrence of two break-points (start and end loci of the event, graphically represented by different shapes in the Figure 1). The events from different nodes (different copy number events) are allowed to overlap. The candidate breakpoint loci are allowed to repeat as a start or end loci of different events. In this way, our model follows the finite sites assumption. In contrast to the infinite sites assumption, which is often made by models operating on single nucleotide variants, the finite sites assumption is much more realistic for copy number events, as the same breakpoint loci with increased genomic instability were observed to be re-used in tumor evolution [51]. The evolutionary tree structure, together with the cells’ attachment to its vertices, defines the copy number events history of each cell and consequently, the set of breakpoints that should be observed within each cell sequence.

We further model the per-breakpoint data assuming that the corrected count absolute differences follow different distributions for breakpoint or no-breakpoint loci, i.e. for loci with or without copy number change in adjacent bins, respectively. Specifically, the corrected count absolute differences in no-breakpoint loci are expected to be close to zero i.e. follow the right half of the normal distribution with th expected value equal to zero and unknown variance depending on the quality of the data. In contrast, for breakpoint loci, the corrected count absolute differences should be close to or greater than one (depending on the magnitude of copy number change) and follow a mixture of normal distributions with unknown expected values and variances. The components of the mixture are expected to correspond to the copy number differences that occur in the analyzed cells at the breakpoint loci. The parameters of those two distributions constitute the set of model parameters. We devise an efficient Markov Chain Monte Carlo (MCMC) sampling scheme that allows to jointly search the vast space of possible CONETs and sample model parameters with cell attachment marginalisation. Summing over the cell attachment reduces the complexity of likelihood calculation. To regularize the model performance for noisy biological data, we employ a set of priors that help obtaining reliable results with desirable tree complexity. We also introduce a count discrepancy penalty that includes the per-bin data and penalizes the model for the inconsistency of corrected counts in bins with the same copy number change history.

After finishing the MCMC sampling step and obtaining the final CONET, we use Maximum Likelihood to attach each of the single cells to one of the tree vertices (Figure 1 C). The final step of the procedure is CN calling, where we further utilize the fact that we can recreate copy number change history for each bin in each cell by reading the path from the vertex to which each cell is attached in the CONET to the tree root. By performing this task for all cells, we define clusters of bins that underwent exactly the same copy number changes thus should have the same integer copy number. We calculate the inferred CN in each bin in each cell by averaging and rounding the corrected counts in each cluster. For all the bins not included in any copy number event, we assume a neutral copy number, thus arriving at the estimated integer CN matrix (Figure 1 D).

### Performance of CONET model on simulated data

To assess the performance of CONET in a setting where the ground truth is known, we conduct the evaluation on simulated data. To this end, we use the generative CONET model, which samples a tree of a given size with attached predefined number of cells and outputs per-breakpoint data in the form of an absolute count difference matrix, according to the assumed absolute count difference distributions (Simulated data).

The difference matrix is used as an input for CONET inference procedure and the results are compared with the ground truth to assess the quality of the output tree structure and breakpoints in each cell, according to cell attachment which maximizes likelihood (ML cell attachment).

Each evaluation scenario is described by three parameters - the size of the CONET tree (with values from {20, 40}), the number of cells (with values from {200, 400, 1000, 2000}) which are randomly assigned to tree vertices, and finally absolute count difference distributions which will be used for the generation of the difference matrices. The number of potential breakpoint loci is fixed to twice the tree size.

Entries of the difference matrix are sampled from two corrected counts absolute difference distributions settings - *well separated* and *poorly separated*. The first one provides a clear distinction between the distribution of the absolute count differences at breakpoint loci and the differences at loci without breakpoints. The well separated setting corresponds to higher quality data, with less noise and higher coverage. The poorly separated setting represents input data with more noise and as such is expected to be more challenging for our algorithm. During the inference procedure, the distributions of the corrected count absolute differences are assumed to be unknown and must be inferred by our algorithm.

Additionally, we run our algorithm with two different choices of priors. The *data size prior* penalizes inference of large trees. The impact of this penalty depends on the constant which is set by the user, and we evaluate this prior with the constant set either to 0.1 or to 0.5. The *attachment prior* encourages our scheme to propose trees that attach cells to nodes with a history consisting of shorter events.

For each of the scenarios described above (number of cells, tree size, distributions setting, prior) we generate 10 random models. For each of the generated models, we run our inference scheme 10 times (each time with a different seed) with 5 ·105 steps for parameter inference and 106 MCMC steps, obtaining 10 inferred CONETs and 10 breakpoint matrices (information about breakpoints in each cell according to their maximum attachment to the tree). The average running time of one inference procedure ranges from less than two minutes in the least computationally demanding scenario (tree size 20 with 200 cells) to half an hour in the most demanding scenario (tree size 40 with 2000 cells). We then compare the inferred data to the ground truth information from the simulated models. This strategy allows us to evaluate not only the quality of a single prediction, but also the consistency of the results across different runs of the algorithm for common input data.

To facilitate the assessment of the quality of inference results we introduce six scores – *Inferred Tree Size, Edge Sensitivity, Edge Precision* and *False Positive Rate, False Negative rate, Symmetric Distance*. Their precise description can be found in section Model evaluation methods for simulated data. Here we only remark that the first three scores assess the quality of the inferred CN event tree. Value of *Inferred Tree Size* should ideally be as close to the size of the real tree as possible. *Edge Sensitivity* and *Edge Precision* quantify the similarity of the inferred tree’s edge set to that of the real tree with their larger values indicating closer similarity. The last three scores assess the quality of breakpoint detection with lower values indicating better inference results. *False positive rate* is the fraction of inferred breakpoints that are not present in the cells according to the true tree and cell attachment, *False negative rate* - a fraction of breakpoints that should be present in cells according to the true tree and cell attachment but are not according to inference results and *Symmetric Distance* measures the average number of incorrectly inferred breakpoints per cell.

Figure 2 depicts aggregated tree scores for the analyzed simulation scenarios and demonstrates the high performance of the model in terms of inferring CONET structure. The sizes of the inferred trees oscillate around the real values of 20 or 40, with slight over- or under-prediction being dependent on the choice of prior (Figure 2 A–D). There is a tendency, however, that the trees grow as the number of cells increases. This is to be expected, since for a higher amount of cells the algorithm has more possibilities of increasing the likelihood by adding subtrees that correspond to small subpopulations of cells. All evaluated priors regularize the model and are effective at limiting the inferred tree growth. For the data size prior, the results depend on the constant controlling the prior’s strength. The more complex scenarios with more cells and bigger true trees, the strong data size prior (constant 0.5) sometimes over-penalizes the tree size, resulting in overly small trees. In comparison to the strong data size prior, the moderate data size prior (constant equal to 0.1) results in trees of size closer to the true values. The attachment prior works in the most subtle way i.e. the model returns the biggest trees compared to the two other priors.

**Figure 2.**
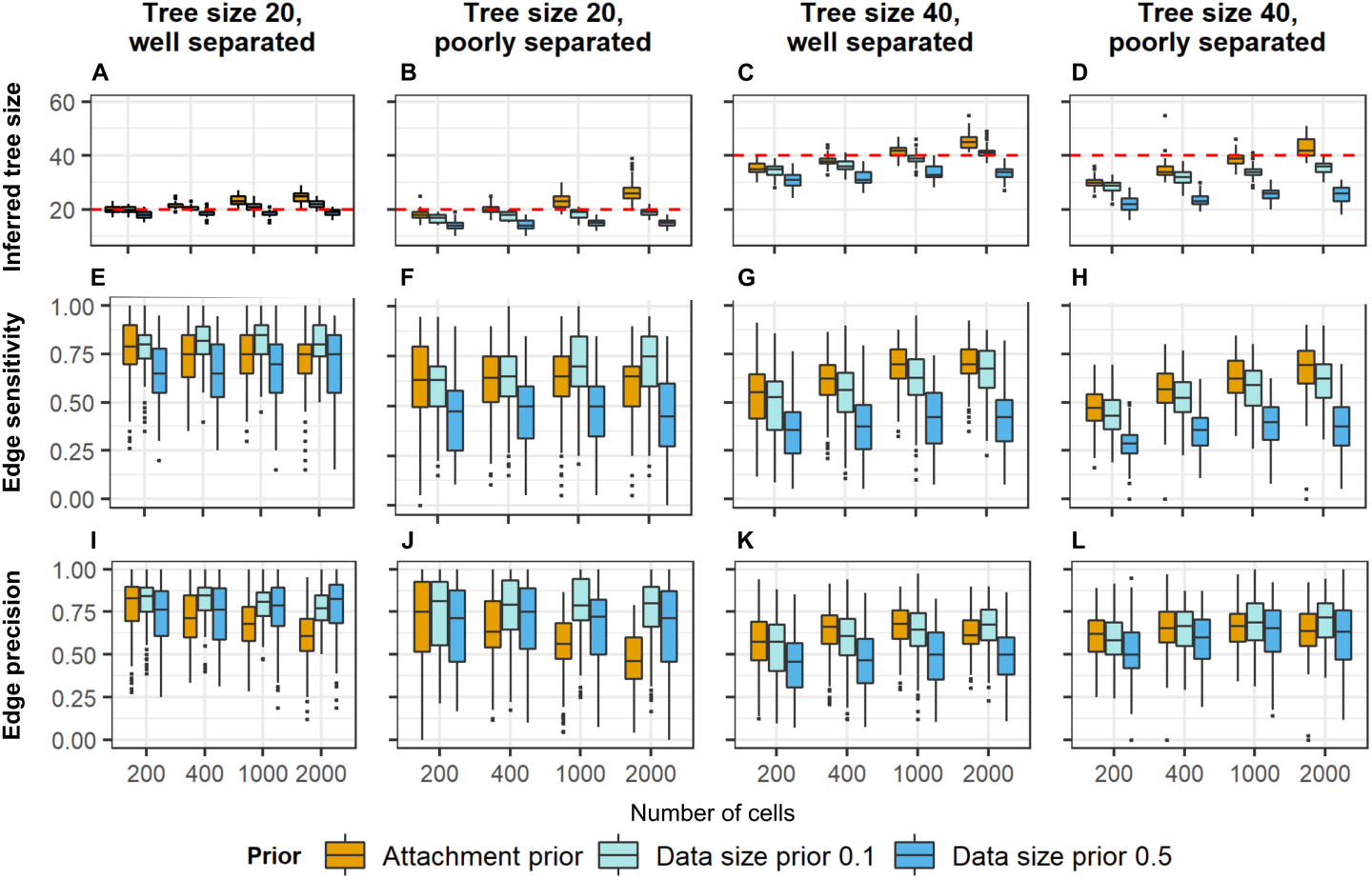
Assessment of the tree structure inference for simulated data tests results. **A-D** Distribution of inferred tree sizes (y axis) depending on the cell number (x axis) for all simulation scenarios. The horizontal line indicates the size of the true event tree. **E-H** Distribution of edge sensitivity (y axis) depending on the cell number (x axis) for all scenarios. **I-L** Distribution of edge precision (y axis) depending on the cell number (x axis) for all scenarios. The results indicate very high efficiency in detecting real event history.

Edge sensitivity (Figure 2 E–H) decreases with stronger regularization due to smaller tree size, but is very high for adequate regularization choice. In simulation scenarios, the strong data size prior gives the smallest edge sensitivity. For trees of size 20 and the case of the well separated distributions (Figure 2 E), both the attachment prior and moderate data size prior yield a high edge sensitivity of around 0.75, regardless of the number of cells. For the same tree size and the poorly separated case, edge sensitivity decreases by around 0.1 for these priors (Figure 2 F). For larger trees (Figure 2 G,H) edge sensitivity further decreases, but it grows with larger numbers of cells. In the most difficult of the analyzed simulation scenarios (tree size 40 and poorly separated data), with 2000 cells, the attachment and moderate data size prior give edge sensitivity of around 0.65.

Compared to edge sensitivity, the different regularization priors have less effect on edge precision (Figure 2 I-L). In all analyzed simulation scenarios, the majority of discovered edges is contained in the real history. In the simplest scenario (tree size 20 and well separated data, Figure 2 I), edge precision is around 0.75 for all cell numbers. Even in the most difficult scenario (Figure 2 L), the median edge precision of the algorithm exceeds 0.5, regardless of the prior and the cell count. In some simulation scenarios (Figure 2 I,J), edge precision decreases with the increasing number of cells for the attachment prior. This is because with this prior, larger trees are inferred.

Figure 3 illustrates the excellent performance of breakpoint detection using our algorithm. In all analyzed scenarios, the median false positive rate of the detected breakpoints is very low. Indeed, regardless of the choice of the prior and the number of cells, the median false positive rate is below 0.1 (Figure 3 A–D). In the well separated data scenarios (Figure 3 A,C), the false positive rates are even lower (median less than 0.05).

**Figure 3.**
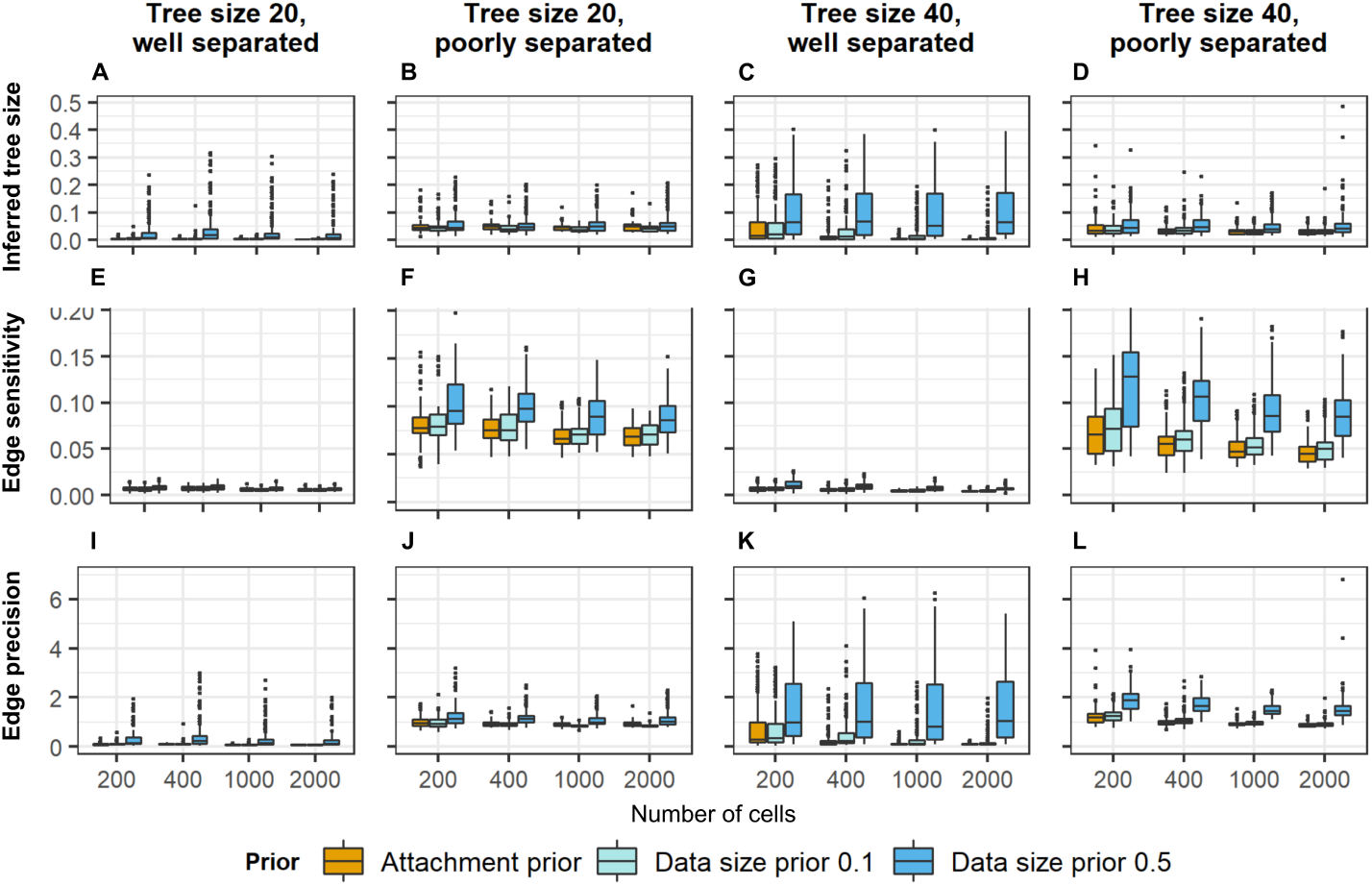
Assessment of breakpoints inference for simulated data tests results. **A-D** Distribution of false positive rate (y axis) as a function of cell count (x axis) for all simulation scenarios. **E-H** Distribution of false negative rat (y axis) as a function of cell count (x axis) for all scenarios. **I-L** Distribution of symmetric distance score between inferred and real breakpoints (y axis) as a function of cell count (y axis) for all scenarios. Overall results are very good for all scenarios, and good data separability is a determining factor of the quality of the results.

Similarly, the false negative rate is very low in all scenarios (Figure 3 E–H). Only in the most difficult simulation scenario (tree size 40 and poorly separated data) and only for the strong data size prior (constant 0.5), the median false negative rate exceeds 0.1. Better data separability (Figure 3 E,G) yields even better results, with median false negative rates below 0.025. Transition to poorly separated distribution results in a more significant deterioration of the false negative rate than of the false positive rate – this is an outcome of more pronounced underestimation of tree size in those scenarios (Figure 2).

Interestingly, both false positive and false negative breakpoint rates are very small for the attachment prior, also in the scenarios with small trees, where attachment prior yielded relatively low edge precision (Figure 2 I,J) and proposed trees were larger than the true trees (Figure 2 A,B). This result suggests that even though edge precision may be worse for CONET with priors that propose trees that are too large, the algorithm can still exhibit good breakpoint detection.

The excellent performance in breakpoint re-detection for simulated data is best quantified using the symmetric difference metric (Figure 3 I–L). Again, even in the most difficult scenario (Figure 3 L), the symmetric distance between real and the inferred breakpoint matrices is only around 1. This means that on average every cell has only a single missed or wrongly inferred breakpoint.

In conclusion, although the best results are achieved for the smaller trees with well separated absolute count difference distributions, overall the algorithm excels in breakpoint detection across all evaluated measures, while the correct choice of regularization ensures satisfactory edge sensitivity and precision.

### Application of CONET to scDNA-seq data of SA501X3F xenograft breast cancer sample

Here we present the application of CONET model to scDNA-seq data from 260 xenograft breast cancer cells in the SA501X3F data set [45], sequenced using the Direct Library Preparation (DLP) method. This specific scDNA-seq technology omits the preamplification step, thus ensuring better coverage uniformity which in turn allows for more reliable CNAs detection.

First, we demonstrate the reasoning behind the choice of regularization used for the inference procedure. Second, we present the inferred CONET, together with breast cancer genes affected by the copy number events on the tree, and the information about the number of cells attached to each vertex. Finally, we present the model-derived CN calls and compare them to the ones published together with the sample in [45], which were inferred using HMMcopy [52] without the aid of joint CN evolution reconstruction.

#### Model calibration for scDNA-seq data sample

In the case of the true scDNA-seq data set, it is advantageous to apply additional regularisation in the form of the count discrepancy penalty for biological data with per-bin corrected count data available. The count discrepancy penalty consists of two terms (Section Definition of the count discrepancy penalty). The first term corresponds to the *L*_2_ distance between the noisy counts in the data and the CN estimation based on the model. The weight of this term is controlled by a constant *s*_1_, with *s*_1_ = 0 corresponding to the fact that this term is not included in the penalty. The second term penalizes trees that create regions changed by CN events and inferred copy number equal to two. The weight of this term is controlled by the constant *s*_2_.

In the first scenario, similarly to the tests on simulated data, we do not apply the count discrepancy penalty at all (*s*_1_ = *s*_2_ = 0). In the second scenario we apply only the first penalty (*s*_1_ = 200000, *s*_2_ = 0). Finally, in the third scenario we take advantage of both count discrepancy penalties (*s*_1_ = *s*_2_ = 200000). In each case, CONET inference is performed under the same MCMC sampling setting (MCMC Sampling): 0.5 million iterations with joint inference of the tree structure (Tree moves) and the corrected count absolute difference distributions (Moves on parameter space), proceeded with 1 million iterations with only tree moves.

The run time for the first scenario is 2.5 hours, the second scenario - 11.5 hours, and the third - just under 10 hours (on a high performance computer with AMD Ryzen Threadripper 3990X 64-Core CPU and 128 GB RAM, using 5 threads). For each scenario, the run times for the whole inference procedures in the case of biological data are longer than for simulated data, since we deal with over ten times more potential breakpoints (which translates to around 100 times more potential CN events). The first scenario is substantially the least computationally demanding because it skips the count discrepancy calculation in each MCMC step and - as a result - infers much smaller trees (tens vs hundreds of vertices). The second scenario runs longer than the third because the inferred trees are larger when we do not penalize trees for inferring regions with CN equal to two.

The obtained CONETs are evaluated using several quality measures (Section Model evaluation methods for biological data). Apart from the tree size (the number of CN events inferred for a given data set), we evaluate the number and the quality of bin clusters that can be defined for each inferred model. Given a CONET, clusters are defined as such sets of those bins across cells that share the same event history, determined by the tree and the cell attachment (Methods). For each model, apart from computing the number of clusters, we evaluate the average cluster support (the percentage of bins in the cluster that have corrected count close to the cluster’s inferred CN, averaged over all bins in the cluster). In addition, we compute the percentage of good clusters (the proportion of clusters with support higher than 70%). The higher the value of the average cluster support and the percentage of good clusters, the better the quality of the inferred CONET. Finally, we evaluate the percentage of clusters with inferred copy number equal to 2. The high value of this parameter indicates that the reconstructed history of CN events is less credible since it is less likely that after series of CN changing events the resulting copy number will return to two.

The analyzed scenarios and proposed quality measures facilitate the comparison of the CONET inference results with and without the additional count discrepancy (Table 1). In the first scenario, the tree inferred without the penalty contains only 35 events and defines 127 clusters. It is hard to expect that such a small tree can fully explain the copy number variation in 260 single tumor cells. Correspondingly, according to all quality measures, this tree has the lowest quality of clusters, compared to the trees obtained with the additional count discrepancy penalty. The tree obtained in the second scenario is noticeably larger, and although the cluster qualities are comparable to the third scenario, the percentage of clusters with CN equal to two is too high. The comparison clearly demonstrates the overall advantage of applying the full count discrepancy penalty (scenario 3), which yields the CONET of the average size and combines the good quality of nodes with low percentage of clusters that infer copy number equal to 2. The fact that the inference without the additional penalty (the first scenario) gives very good performance on simulated data, shows that the real biological data poses a significantly more difficult challenge for the model.

**Table 1.** Assessment of CONET inference for biological data.

| Scenario | Tree size | No of clusters | Avg cluster support | Perc of good clusters | Perc of CN=2 clusters |
| --- | --- | --- | --- | --- | --- |
| 1 | 35 | 127 | 83% | 69% | 17% |
| 2 | 380 | 3136 | 89% | 89% | 16% |
| 3 | 225 | 905 | 90% | 90% | 4% |

Second, we evaluate the models obtained in the three scenarios with respect to the quality of CN estimation (Methods; Table 2). As above, the evaluation is based on the quality of bin clusters (sets of bins that share the same event history according to the model). Here, we first consider bin clusters that are indeed changed by events of the tree, referred to as *in events*. Second, we consider bins that are in one large cluster of all bins that were not included in the events of the tree, i.e. the tree does not infer any copy number tree for these bins. These bins are referred to as *outside events*. To quantify the dispersion of corrected counts in both the bin clusters inside and outside events, we use the Gini index and Shanon Entropy measures. Next, copy number true positive rate (CN-TPR) is computed as the fraction of bins with true corrected count not rounding to two and contained in events of a CONET, out of all bins with true corrected count not rounding to two. Here, we consider the bins with true corrected count not rounding to two as the positive examples of bins that truly underwent a copy number change event. The copy number false positive rate (CN-FPR) quantifies the fraction of bins that are part of a CONET events (the bins that underwent a CN event according to our model) out of the total number of bins with corrected count rounding to two (the bins that did not undergo a CN event according to the data). Root mean square error for our CN calling procedure (CN-RMSE) is calculated as the quadratic mean of the differences between inferred integer CN and corrected count in each bin, i.e. the square root of the count discrepancy penalty. For a correctly inferred CONET, the CN-RMSE reflects the noisiness of scDNA-seq data.

**Table 2.** Assessment of CN calling for biological data.

| Scenario | Avg<br>Gini Index<br>in events | Avg<br>Entropy<br>in events | Gini Index<br>outside<br>events | Entropy<br>outside<br>events | CN-TPR | CN-FPR | CN-RMSE |
| --- | --- | --- | --- | --- | --- | --- | --- |
| 1 | 0.12 | 0.43 | 0.22 | 0.57 | 27% | 20% | 0.74 |
| 2 | 0.08 | 0.20 | 0.10 | 0.34 | 96% | 73% | 0.37 |
| 3 | 0.09 | 0.16 | 0.11 | 0.37 | 84% | 9% | 0.40 |

The small size of the CONET inferred without the count discrepancy penalty in the first scenario, results in bad overall quality of inferred CNs according to all quality measures. Compared to the trees inferred with the penalty, this tree has higher dispersion inside events and - more apparently - outside events, very low CN-TPR and almost two times higher CN-RMSE. In the second scenario an opposite situation occurs, where the CN-TPR is close to 100% because this most complicated CONET includes such a high number of events that almost all bins with CN far from two are included inside them. The dispersion measures improve substantially compared to scenario 1 and the CN-RMSE is the lowest. This happens at the cost of CN-FPR, which is over three times higher than in the first scenario and nine times higher than in the third scenario. This again proves that without penalizing inference of copy number equal to two inside events, the inferred tree grows too large. The CN calling results obtained using the CONET with the full discrepancy penalty in the third scenario are by far the best, constituting a well balanced compromise between the too simple and too complicated CONETs from the two other scenarios. The quality of CN calling results for the tree in scenario 3 is higher than for the first scenario across all parameters. In comparison to the tree obtained in scenario 2, this tree is similar in terms of low dispersion in counts in bins inside and outside events, rare inclusion of bins with corrected count far from two in CN events (CN-TPR) and low CN calling error (CN-RMSE), while scoring far better in terms of restricting the inclusion of bins with corrected count close to two (CN-FPR).

We conclude that additional regularisation in the form of the full count discrepancy penalty is necessary when dealing with noisy, low-depth scDNA-seq data from real experiments. The degree of this regularisation should be calibrated to the specific biological data set with scDNA-seq technology in mind, utilising different *s*_1_ and *s*_2_ count discrepancy constants’ values and comparing the results with the aid of different quality measures described in Model evaluation methods for biological data.

#### CONET for SA501X3F xenograft breast cancer data

Figure 4 A presents the CONET inferred using the full count discrepancy penalty described in the third scenario from the previous section. The complex structure of the obtained CONET illustrates the high level of instability in the cancerous genome. In total, the events in the tree overlap with 27 genes determined as associated with breast cancer by the COSMIC Cancer Gene Census [53]). Additional file 1 presents the comprehensive list of all the CONET vertices with information about their parent, genomic coordinates of each event and breast cancer genes concerned by the event.

**Figure 4.**
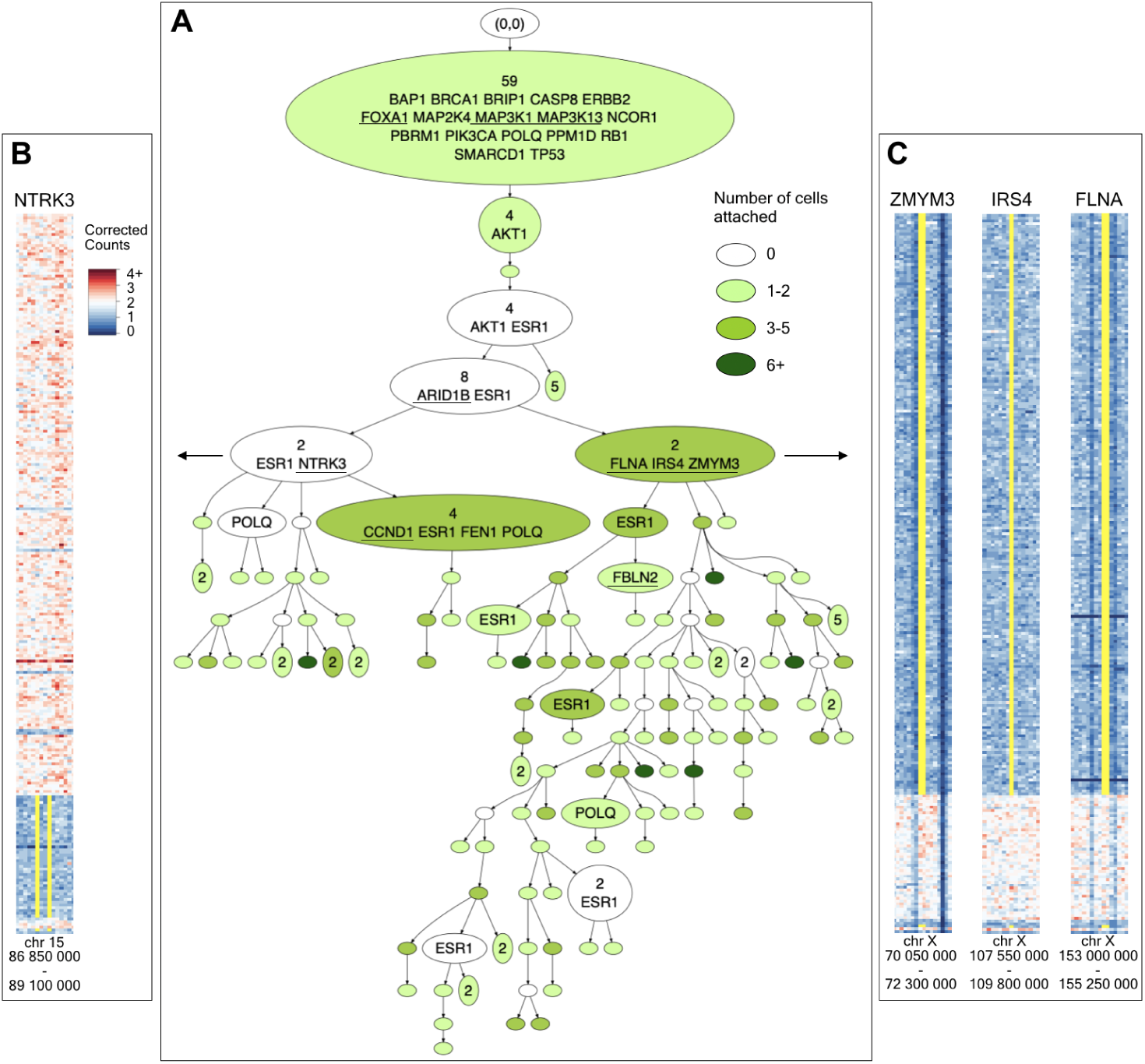
CONET for SA501X3F xenograft breast cancer data set. **A** The CONET is drawn in a compacted form, with part of vertices collapsed when it is possible without losing important information, i. e. when the collapsed vertex has no cells attached and not more than one child. Specifically, the collapsed vertex is joined with its closest descendant that does not satisfy these criteria (number of events is shown at the beginning of joint vertices). This results in decreasing the tree size from 225 to 131. The number of cells attached to each vertex is illustrated with different colors, where white vertices have no cells attached and the darker green indicates more cells attached. The most biologically significant information are the names of the breast cancer genes [53] concerned by copy number events, which are printed in alphabetical order in all internal vertices. Underlined genes appear only once in the CONET. **B, C** Enlarged parts of corrected counts matrix heatmap are visible on the sides with areas around breast cancer genes that divide the cells into two distinctive subpopulations. The yellow lines are the borders of each of the genes drawn for cells that have copy number change of the gene according to their attachment in the CONET.

Looking closer at the inferred CN events history, we first observe a linear evolution of the tumor, where the vast majority of cells undergo a series of common CN events, visible in the trunk of the tree. Those events overlap with many characteristic breast tumor suppressor genes such as *BRCA1, TP53, RB1* or *CASP8* [53]. Copy number events that can cause a decrease in expression of those genes (all genes fall into clusters with inferred CN equal to one - see Figure 5) could have contributed to the onset of breast cancer. The next significant event in the evolutionary history of this tumor sample is the branching of the trunk that divides the cells into two distinct subgroups, the smaller one (left of the tree in Figure 4 A) characterized by unique NTRK3 gene copy number change, and the more abundant (on the right of the tree in Figure 4 A), distinguished by unique *FLNA, IRS4* and *ZMYM3* copy number changes (with inferred copy number equal to one). Figure 4 B presents an enlarged fragment of corrected counts heatmap with the genomic region around *NTRK3* where the gene location and subpopulation of cells with lowered CN according to the inferred CONET is marked in yellow. The second subpopulation of cells which emerged during main tree branching is shown in Figure 4 C across fragments of corrected counts heatmaps surrounding *FLNA, IRS4* and *ZMYM3*. It can be speculated that the copy number event concerning the *NTRK3* gene could have been part of a known translocation causing the formation of the *ETV6-NTRK3* fusion oncoprotein characteristic of human secretory breast carcinoma [54]. This event could result in an evolutionary advantage for the subpopulation of cells that are attached under the vertex describing this copy number event. Among the genes characterising the other larger subpopulation, *ZMYM3* is a known tumor suppressor [53], whose deficiency impairs DNA repair by homologous recombination and can result in high genome instability [55]. Evidence of this instability is clearly visible in the CONET structure for this subpopulation where we distinguish numerous CN events that do not overlap with new breast cancer genes. Another interesting observation can be drawn from genes such as *ESR1* or *POLQ* (both falling into regions with amplified copy number for most of the cells - not shown) that reappear many times in the events of the inferred CONET, also in parallel branches, suggesting that some genomic regions have very high instability and are more prone to CNAs. This also points to the fact that the evolutionary trees reconstructing copy number changes do not follow the perfect phylogeny assumption.

**Figure 5.**
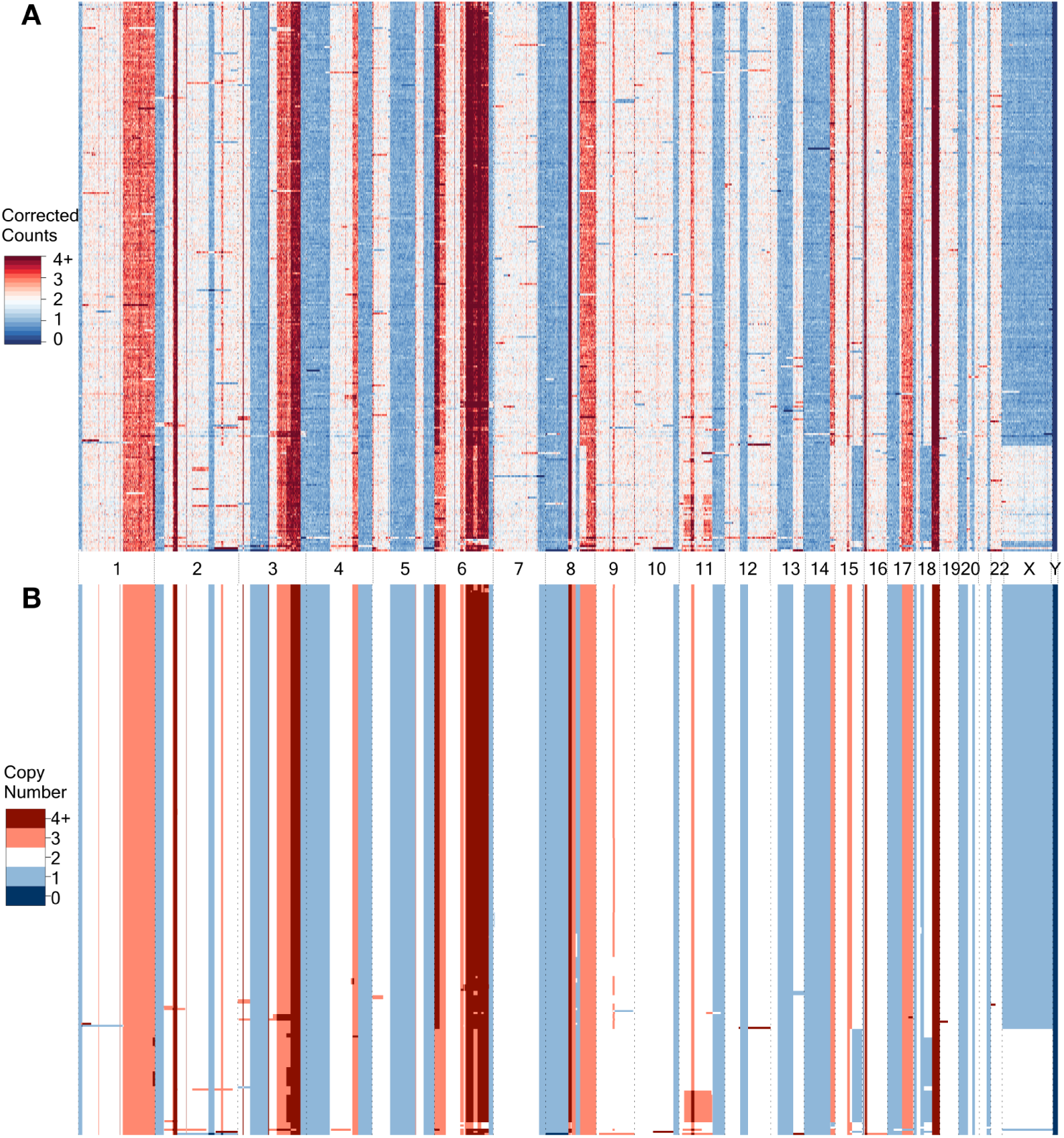
Graphical illustration of CN calling procedure for SA501X3F xenograft breast cancer data set. **A** The CC heatmap illustrates the biological data with corrected counts in genomic bins, **B** the CN heatmap presents the inferred integer CN for equivalent bins. Columns correspond to genomic locations, rows to single cells. The rows in both matrices are in the same order fixed using hierarchical clustering of cells according to their inferred copy numbers.

#### Copy number calling for SA501X3F xenograft breast cancer data

Figure 5 graphically represents the high quality of the CN calling procedure for SA501X3F xenograft breast cancer data. The regions with clearly visible deviations from the neutral copy number in the corrected counts heatmap (Figure 5 A) are for the most part correctly identified as such in the inferred integer CN heatmap (Figure 5 B). At the same time, the inferred CN heatmap lacks the noise observed in the true corrected count data. Importantly, the fidelity of CN events’ reconstruction is maintained even for very narrow genomic events (sometimes up to one bin wide). This is especially true for CN events that are common for a large enough number of cells.

#### Comparison to other methods

To demonstrate the advantage of the CN calling procedure, we compare it to HMM-copy [52], the CN calling method developed specifically for this type of scDNA-seq data. We run HMMcopy using the same SA501X3F xenograft breast cancer data. The inferred HMMcopy CN matrix is used to calculate CN-RMSE the same way as was done for our CN calling procedure results. The CN-RMSE for HMM is equal to 0.75, which is similar to CN-RMSE obtained by the CONET in the first scenario described in Model calibration for scDNA-seq data sample and two times higher than the best result of the CONET described in the second scenario. This clearly demonstrates that using CONET to call integer copy number in genomic bins improves the fitting of inferred CN matrix to the real data compared to the method that does not involve joint inference of CN evolution.

## Conclusions and discussion

Recent developments of high-throughput scDNA-seq technologies have enabled tracing copy number changes in single cell genomes. In this work, we introduce CONET, a novel Bayesian model for inference of an evolutionary tree of copy number events and copy number calling for scDNA-seq data. CONET is robust to errors in scDNA-seq data, as it models the per-breakpoint and per-bin readouts in a probabilistic manner. By combining the process of tree inference and copy number calling, it gains power in both tasks. We propose an efficient MCMC procedure for the search across the space of possible trees and model parameters.

CONET performs favorably in terms of evolutionary tree reconstruction and copy number calling both on simulated and real data. Comprehensive analysis on simulated data indicates that CONET excels in the correct breakpoint detection for single cells, even in the case of more demanding scenarios with bigger trees and more noisy data. Appropriate choice of regularization priors enables accurate inference of the model structure, as measured by the edge precision and sensitivity.

Using CONET, we analyzed the evolutionary history of copy number events and copy number changes in single cells for a xenograft breast cancer sample. Evaluation of the trees obtained using different regularization schemes indicates that in the case of modeling real data, penalization for the disagreement between the estimated and actual per-bin counts is crucial for constraining the model and obtaining high-quality trees. Comparison of the corrected per-bin data with inferred copy number profiles illustrates the excellent performance of breakpoint detection and the CN calling procedure for biological data. We observe the smoothing of the noisy biological signal for less evident events. Since the biological truth is unknown, it is difficult to assess the quality of the inferred CONET structure. Still, the presence of many genes important for breast cancer evolution in the trunk of the inferred tree and division of cells into two distinct subclones, testifies to the fact that our model can be of help in deconvoluting evolutionary relationships between cancerous cells and identifying driver events.

To our knowledge, CONET is the first Bayesian probabilistic approach for copy number evolution inference and copy number calling, that fully exploits the scDNA-seq readouts, in the form of both per-breakpoint and the per-bin data. CONET differs from other recent evolutionary models of breakpoints or copy number events: the model of [48], MEDALT [49] and SCICoNE [50]. For the trees inferred by [48] or MEDALT, the nodes do not correspond to copy number events. Each node of [48] tree corresponds to acquisition of only a single breakpoint. The MEDALT model is a lineage tree spanning the input cells. In contrast, SCICoNE explicitly models an event tree, where each node is labeled with a vector of starts and ends of CN events, together with the copy number change that occurred for each of the events. We deliberately avoid modeling the exact copy number changes acquired at each event, thereby vastly reducing the space of possible models.

The reduced space and efficient MCMC implementation facilitate advantageous computation times for CONET inference procedure. The main factors influencing the run time are the number of cells and the size of inferred CONETs, which can be regulated by the user with the number of potential breakpoints, data size priors and the choice of corrected counts penalty. The control over model parameters and regularization will give a particular advantage when dealing with the much larger and hopefully less noisy data sets, which can be anticipated in the future, given the constant advances in whole genome scDNA-seq techniques.

Given that a tumor sample was sequenced using both scDNA-seq with high coverage uniformity and using such other techniques that allow SNV calling, the output from CONET can be utilized to enhance approaches analysing SNV evolution. For example, the accurate CN calls from CONET can provide valuable input to methods that rely on CNA information for improved elucidation of tumor subclones and their relationships from variant allele frequencies in bulk sequencing data [9, 10]. Compared to the CN calls alone, CONET output can define even more specific constraints for reliable modeling of evolutionary SNV trees inferred from high coverage scDNA-seq [56].

Deciphering the gene copy number evolution of tumors is of huge importance for the understanding of the process of carcinogenesis. Application of CONET to scDNA-seq data from tumor samples of patients enables identification of copy number events that occur early in tumor evolution, amplifying or deleting cancer driver genes in the entire population of tumor cells. For the SA501X3F breast cancer sample, the trunk genes included *BRCA1, TP53, RB1* or *CASP8*, all of which play pivotal roles in breast cancer progression. Such trunk alterations could potentially suggest the choice of efficient anticancer treatment. On top of that, our approach can identify distinct subclones characterised by unique gene copy number alterations caused by events from specific tree sub-branches. For the breast cancer application, examples of such genes identified as subclonally altered are *NTRK3* and *ZMYM3*. The alterations of these genes can in turn confer resistance to therapy, and as such can highlight the need for additional therapeutic intervention or combinatorial treatment.

In summary, CONET is a powerful tool that takes advantage of scDNA-seq to better understand the copy number evolution in cancer and may guide the choice of therapy when applied to patient data in the clinic. Ultimately, once scDNA-seq data from larger cohorts is available, the application of CONET will help to uncover general patterns of the order of occurrence of CNAs in tumors together with their importance.

## Methods

### Real data preprocessing

To illustrate the performance of our model on true biological data we apply it to scDNA-seq data from 260 xenograft breast cancer cells SA501X3F data set [45] sequenced using Direct Library Preparation (DLP) method. According to [45] the sequencing reads were binned into 150 kilo base bins, corrected for GC content and mappability and normalized such that 2 signifies a neutral copy number. The resulting positive real numbers for each bin are further called corrected counts in bins and constitute the per-bin input to our model. We calculate the corrected count absolute differences by subtracting corrected counts in the adjacent bins and taking the absolute value. The resulting corrected count absolute differences matrix with genomic loci in columns and cell ids in rows represents the per-breakpoint input to the model.

To run our model, we also need to establish candidate breakpoint loci set. The set of candidate loci determines the set of possible copy number events and as a consequence significantly influences the computational complexity of the model inference procedure. The candidate breakpoints are set by the user.

Here, we run HMMcopy [52] implemented as Bioconductor package using corrected counts as input with standard parameters, obtaining integer copy number state for each bin. We further assume that each locus with inferred copy number state change in adjacent bins is a possible breakpoint. To this set of candidate breakpoints we also add the beginning and end of each chromosome, together with loci which show high corrected count absolute difference evidence (more than 80% of cells with the corrected count absolute difference higher than 3). To complete our potential breakpoint loci set, for each of the above mentioned candidates (except chromosome ends) we add one locus to the right to be always able to infer short one bin events. The final set of candidate loci for the SA501X3F data set contains 2044 possible breakpoints. After this step, we prepare the final corrected count absolute differences input matrix restricted to chosen candidate breakpoint loci columns.

### CONET - Copy Number Event Tree model

Below, we explain CONET, a generative probabilistic model for inferring tumor evolution on single cell copy number events. Let 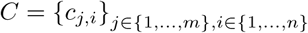 denote a data set of corrected read counts from *n* genomic bins for a total of *m* cells. We assume the read count data are pre-processed, in particular, normalized to neutral copy number 2, corrected for GC content and other potential biases. For each chromosome we add an artificial bin representing the end of a chromosome with corrected count equal to 2. This is necessary to being able to include copy number events that start or end at the ends of a chromosome, i.e., physically have only one breakpoint. The data modeled by CONET is defined as the absolute differences of counts at consecutive bins:

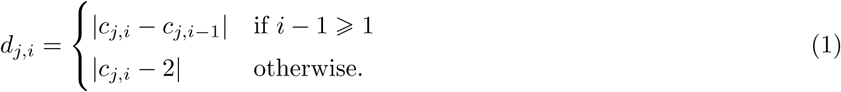

Each difference is indexed with *j*, denoting the cell it is calculated for, and with *i*, denoting the genomic locus (or interchangeably - the genomic bin starting in this locus). We assume a set of loci *L* ⊂ {1, …, *n*} is given. We define a copy number event as an ordered pair of loci (*i, l*) from *L*, such that *i < l*, i.e., locus *i* occurs before *l* in the genome, and loci represented by *i, l* lie on the same chromosome. We assume that in bins [*i, i* + 1, …, *l*) the copy number changed during the evolutionary history of cells that underwent this event. Since the loci *i* and *l* mark the endpoints of the deleted or amplified regions, we refer to them as breakpoints. Loci from *L* will be referred to as candidate breakpoints.

We assume that each copy number event can occur only once in the tumor evolution. Still, one breakpoint can be part of many copy number events and the events can overlap. In this sense, our tree does not satisfy the infinite sites assumption.

Let *D* denote the data matrix of absolute count differences for such bins that start at candidate breakpoints, i.e., such that for *d*_*j,i*_ it holds *i* ∈ *L* and *j* ∈ {1, *…, m*}.

CONET is a tuple (*T, σ, θ*), where *T* is the evolutionary tree structure, *σ* is referred as cell attachment, and *θ* denotes the set of model parameters. Let *T* denote a directed rooted tree with a set of vertices *V*_*T*_ corresponding to copy number events and edges to the partial order of these events. Additionally, *V*_*l*_ denotes the set of all the tree leaves and *V*_0_ denotes the set of all possible events, which are not present on the tree (i.e., that do not belong to *V*_*T*_). We refer to *V*_0_ as the set of inactive events, an(*v*) denotes the ordered set of vertices (including vertex *v*) that occur on the path from vertex *v* to tree’s root and depth(*v*) denotes the depth of vertex *v* in the tree. The root of the tree is given as an artificial event denoted by (0, 0), corresponding to no somatic copy number events.

The attachment of cells to the tree is given by a function *σ* : {1, *…, m*} → *V*_*T*_. For a given cell *j*, the path from *σ*(*j*) to the root of the tree defines the set of events that the cell *j* underwent and their order (an(*σ*(*j*))). It also specifies the set of breakpoints that this cell has. Indeed, for a given tree *T* and cell attachment *σ*(*j*), the set of breakpoints of cell *j*, denoted by *I*_*bp*_(*j*|*T, σ*) is given by the breakpoints that occur in at least one event on the path an(*σ*(*j*)). We also define *I*_0_(*j*|*T, σ*) to be the set of remaining candidate breakpoints in *L*, i.e., *I*_0_(*j*|*T, σ*) := *L \ I*_*bp*_(*j*|*T, σ*).

The set of parameters *θ* parametrizes the distributions of the absolute count differences, depending on whether they are calculated at breakpoints or not. The absolute count differences at breakpoints are expected to be large, while differences at loci that are not breakpoints should oscillate around zero. Accordingly, for a given tree *T* and attachment *σ*, we have *d*_*j,i*_ ∼ *f* (*d*_*j,i*_|*T, σ*_*j*_, *θ*), where

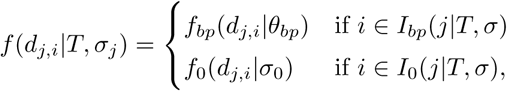

and *f*_*bp*_ is a mixture of *K* normal distributions truncated to ℝ_+_ and *f*_0_(·|*σ*_0_) is a mean zero normal distribution, also truncated to ℝ_+_. Thus, 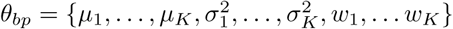, where *µ*_1_, *…, µ*_*K*_, are the means, 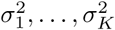 are the variances and *w*_1_, *…, w*_*K*_ are the weights of the mixture components. The full set of CONET parameters is given by *θ* = *θ*_*bp*_ ∪ {*σ*_0_}. The number of mixture components *K* is user defined. During the optimization procedure *K* is decreased when the corresponding weights decrease beyond a user defined threshold.

Thus, the likelihood of the data *D* (the count difference matrix), given unknown model constituents (tree structure, attachment, parameters) is defined to be:

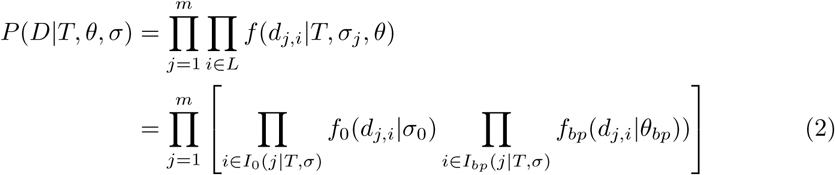

Intuitively, the likelihood is expected to be high for such tree structures and attachments that indicate breakpoints for count differences with relatively high values and vice versa: no breakpoints for small count differences.

We assume that cells’ attachment probabilities are conditionally independent given the tree structure and do not depend on *θ*:

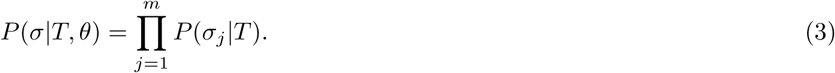

The exact form of *P* (*σ*_*j*_|*T*) is specified below (Section Attachment prior). To decrease the complexity of the state space of our algorithm, we marginalize out the cell attachment vector *σ*. We furthermore use assumption (3) to transform the marginalized version of formula (2) to achieve *O*(|*L*| *· m ·* |*V*_*T*_ |) complexity of a single likelihood computation:

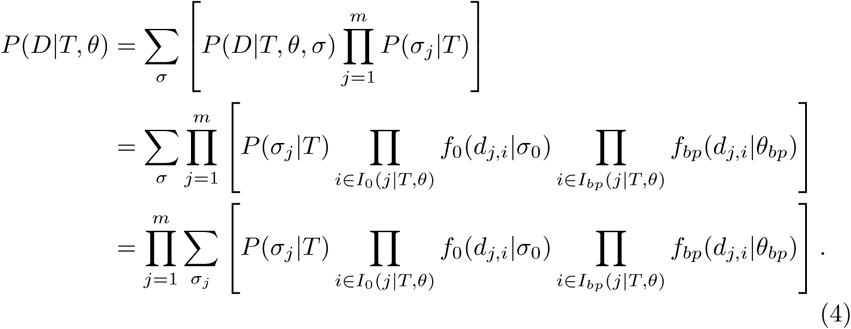

Cell attachment information is obtained from (*T, θ*) by choosing such *σ*, which maximizes *P* (*D*|*T, θ, σ*). Such *σ* is denoted by *σ*\* and referred to as *maximal attachment*. The described generative model deliberately ignores the actual copy number change associated with each event and the actual count data observed at each bin. In this way, the state space for our model is significantly reduced. Still, the copy number state for each event and each bin in each cell can easily be estimated. We utilize this fact to 1) penalize the model for inconsistency between the estimated copy number state and counts in each bin and cell during training and 2) estimate the state in the bins for the final model (see below).

### Priors

Here, we define the priors: *P* (*T*) for tree structure, *P* (*θ*) for the parameters and *P* (*σ*_*i*_|*T*) for attachment given the tree structure. In all formulas below, *k*_*x*_ are model hyperparameters that are set by the user.

#### Tree structure priors

We define the tree structure prior as a product of three different prior distributions, i.e.,

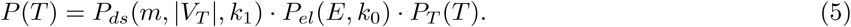

The *P*_*ds*_ is a prior for the tree size, following the Occam’s razor principle, that the simplest explanation is the most probable. The prior *P*_*ds*_ controls the growth of simulated tree size and prevents overfitting of the tree structure and is defined as follows

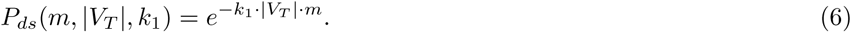

This prior explicitly depends on the number of cells *m*. This constant is used to avoid over-fitting of the model to the data and constrains excessive growth of the tree size with increasing sample size.

The prior *P*_*el*_ accounts for the observation that shorter copy number aberrations occur more often than the long ones [51], and favors trees with smaller total length of copy number events *E*_*T*_, i.e.:

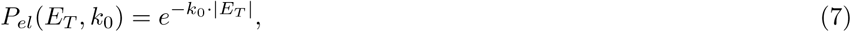

where 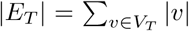 and |*v*| denotes the length of event *v* (the position of the end locus minus the position of the start locus).

The last prior is technical and serves as a stabilizer for proposal ratios of those MCMC sampling moves that change the tree size (Add leaf, Remove leaf).

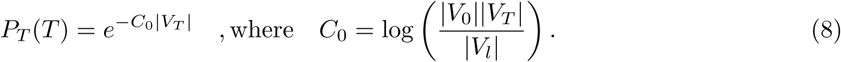

### Attachment prior

The attachment prior *P* (*σ*_*j*_|*T*) reflects our belief about probable attachment of cell *j*. We can choose between an uniform attachment prior or assume that cell attachment depends on the tree structure and the resulting copy number events. In the latter case the prior is proportional to

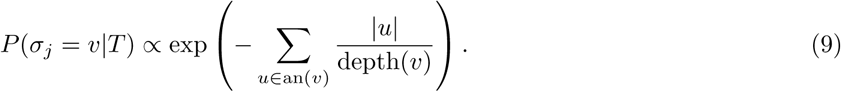

During cell attachment marginalisation the second alternative attributes more weight to attachments that correspond to a shorter average length of events in cells.

### Parameters priors

Recall that the set of parameters consists of 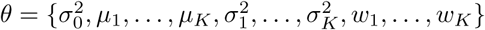. To facilitate MCMC sampling of *θ*, we instead work with the parameters’ set

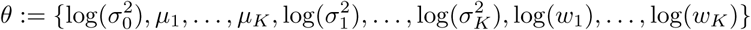

(denoted also by *θ* with a slight abuse of notation). Moreover, we do not force the log-weights to sum to one – this enables moves which change only one of the weights and enhances exploration of state space.

Priors for each of the 3*K* + 1 transformed one-dimensional parameters’ distributions are independent centered normal distributions (for means priors are truncated to ℝ_+_) with variances *ν*_*i*_ : *i* = 1, *…*, 3*K* + 1. In our simulations we set *ν*_*i*_ = 1 for all *i*.

### Count discrepancy penalty

Here, we describe 1) the estimation of the copy number states of bins in each cell given a certain event tree *T* and 2) penalization of the model for the inconsistency between the estimated states and actual counts.

#### Estimation of copy number states for each bin in each cell

Consider all possible bin-cell pairs, denoted (*i, j*), for *i* ∈ {1, *…, n*} and *j* ∈ {1, *…, m*}. We say bin *i* is contained in event *v* = (*l*_1_, *l*_2_), denoted *i* ∈ *v*, if *i* is larger or equal than *l*_1_ and lower than *l*_2_.

For a given tree *T* and cell attachment *σ*, consider such a bin-cell pair (*i, j*) for which there exists an event *v* ∈ an(*σ*(*j*)) that satisfies *i* ∈ *v*, i.e., according to the model, bin *i* has its copy number changed during cell’s *j* evolutionary history. Note that since we allow events in *V*_*T*_ to overlap, there may be several such events in an(*σ*(*j*)). Denote by *v*_*F*_ (*i, j*|*T, σ*) the set of all such events in an(*σ*(*j*)). In the evolutionary history of the tumor described by *T* and *σ, v*_*F*_ (*i, j*|*T, σ*) is the set of all events that affected the copy number state of bin *i* in cell *j*.

For a bin-cell pair (*i, j*) there may be no such vertex *v* ∈ an(*σ*(*j*)) that *i* ∈ *v*. In this case, according to *T* and *σ*, the copy number state of bin *i* in cell *j* was not changed during the evolutionary history of the tumor. Such a bin has the same copy number state as the root (0, 0) of the tree, and we fix *v*_*F*_ (*i, j*|*T, σ*) = ø.

Given an event tree *T* and attachment *σ*, we define a clustering ℬ 𝒞 = ⋃ *BC*_*w*_, where each cluster *BC*_*w*_ is defined by bin-cell pairs with the same *v*_*F*_ (*i, j*|*T, σ*)

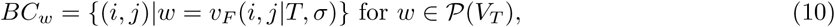

where 𝒫 (*V*_*T*_) denotes the powerset of vertices. For *w* ≠ ø, *BC*_*w*_ is a set of bins that share events that changed their copy number state. In contrast, *BCø* is the set of bins that did not have their copy number state changed.

Note that for (*T, σ*) reflecting the true copy number event history, all corrected counts in bins belonging to a given *BC*_*w*_ should be approximately equal (modulo measurement noise) and reflect the true copy number in those bins. Thus, we estimate the copy number in bins from *BC*_*w*_ using the average count in cluster

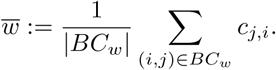

For the root node cluster *BC*_ø_, prior knowledge tells that 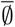 should be equal to 2. (this value can be adjusted to incorporate other scenarios) Thus, in this special case we fix 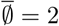.

#### Definition of the count discrepancy penalty

To score how a given event history fits to corrected count data we define

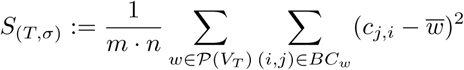

The *S* score can be seen as the L2 distance between noisy counts *C* and the CN calling results of the model.

Furthermore, we define a second score, *S*′ which counts the number of bins for which the average count is close to 2. The penalty *S*′ is motivated by the fact that a bin with copy number 2 is not expected to be a result of copy number evolution. Such bins could result for example from an amplification by one copy and then deletion by another copy, but such a situation, by Occam’s razor, is less likely. Thus, we define

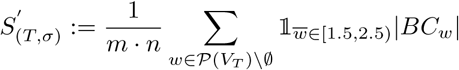

which penalizes events with inferred *CN* = 2.

Finally, we define the count discrepancy penalty as

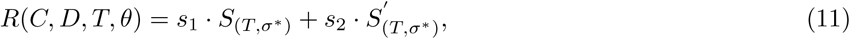

where *s*_1_ and *s*_2_ are user defined non-negative constants and *σ*\* is the maximum likelihood attachment of cells calculated from (*D, T, θ*). In the case when the data *C* is not available, we set *R*(*C, D, T, θ*) = 0. This situation occurs when the data is simulated from the generative model, as CONET only generates the count differences *D*.

### Copy number calling

Given an event tree *T* and the count data, we form the clustering ℬ 𝒞 = ⋃ *BC*_*w*_. The estimated copy number state for each bin in each cell mapped to *BC*_*w*_ is given by the integer number that is closest to *w* with maximum inferred CN equal to 10.

### MCMC Sampling

Our approach is to maximize the penalized *a posteriori* distribution *P* (*T, θ*|*D*). *P* (*T, θ*|*D*) is proportional to *P* (*D*|*T, θ*) *· P* (*θ*) *· P* (*T*) while the penalized version of the log-distribution is equal to:

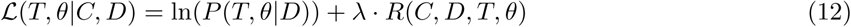

To this end, we employ a standard Metropolis-Hastings (M-H) algorithm on the joint space of event trees and mixture parameters (attachment variables have been marginalized out to decrease the size of the state space).

Moves on the state space are divided into two groups - those that change the count difference distribution parameters *θ* and those that modify the tree structure *T*. We switch between those two types in an alternating fashion. For one parameter update, we perform a fixed number of tree changing moves (in the case of our simulations this number was set to 10). This is motivated by the fact that moves on *θ* are more computationally intensive.

We use *q*(*T* ′, *θ*′|*T, θ*) to denote the proposal kernel. Note that by the discussion above always either *T* ′ = *T* or *θ*′ = *θ*. When it causes no confusion we omit fixed variables from the kernel, so for instance the kernel for the moves that change tree structure will be denoted by *q*(*T* ′|*T*).

Move acceptance probability is standard and given by the expression

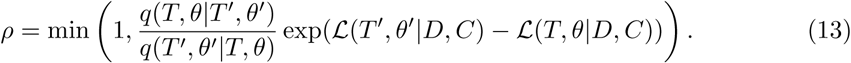

### Tree moves

We employ standard moves on the space of trees, i.e., *Prune and reattach* and it’s combinations: *Swap subtrees* and *Swap vertices on the tree*. Additionally, we employ moves that change *V*_*T*_ : *Add leaf, Remove leaf, Swap event ends between the tree vertices* and *Swap vertices between V*_*T*_ *and V*_0_.

A tree of required size can be obtained from any tree by subsequent applications of *Remove leaf, Add leaf* moves. Structure of a tree can be adjusted by *Prune and reattach* moves. And finally any labeling can be obtained by *Swap vertices between V*_*T*_ *and V*_0_ moves. Hence our tree sampling scheme is irreducible. Aperiodicity of *Prune and reattach* assures aperiodicity of the whole chain. Below we explain the moves in detail.

#### Prune and reattach

We sample a vertex *v* uniformly from the tree and cut the edge leading to this vertex to remove the subtree rooted at *v* from the tree. Then we sample one of the remaining vertices (including the root) uniformly and attach the subtree there instead. The reverse of this move, where we again sample *v* first but then pick its old parent, has the same proposal probability since the non-descendant set has the same size each time *v* is removed. Since we can also choose the old parent when sampling a new one, this move has a non-zero probability of proposing the same tree *T*, ensuring aperiodicity. There is also a path from any tree to a tree with all vertices attached to the root, by moving each vertex to the root step by step. Via reversibility, we can likewise move from there to any other tree.

#### Swap vertices on the tree

We sample two vertices uniformly from the set *V*_*T*_ *\* {(0, 0)}

(we can not relabel the root) and exchange their positions on the tree. To reverse the move, we need to resample the same vertices. Hence, the proposal kernel is symmetric.

#### Swap subtrees

We sample two vertices *u* and *v* uniformly from the set *V*_*T*_ *\* {(0, 0)}. If the two vertices are not in an ancestor/descendant relationship we detach them (together with their subtrees) from their parents and reattach them to each others’ former parent. Since for the reverse move we would simply need to select the same pair of vertices, this case is symmetric.

In the other case, assume *v* is a descendant of *u*. First, we cut the edge leading to *v* and move it with its subtree and attach it to the parent of *u*. Next, we detach *u* and its remaining subtree (with *v* and its descendants removed). The new parent of *u* is sampled uniformly from among *v* and all of its descendants. This assures the move to be reversible. To reverse the move, we again need to sample *u* and *v* at the start, and also to sample the previous parent of *v* from among *u* and its new (remaining) descendants. If we denote the number of descendants of *u* as *d*(*u*), the proposal probabilities now depend also on *d*(*u*) and *d*(*v*), i.e.,

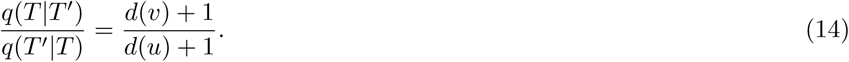

#### Swap vertices between V_T_ and V_0_

We sample a vertex *u* uniformly from the set *V*_*T*_ *\* {(0, 0)} and a vertex *v* from |*V*_0_| available inactive events. Then we transpose those vertices: we put *v* on the tree instead of *u* and move *u* to *V*_0_. To reverse the move, we again sample *v* first from |*V*_*T*_ *\* {(0, 0)}| and *u* from |*V*_0_| available inactive events. The reverse move has the same proposal probability since both *V*_*T*_ and *V*_0_ have the same cardinalities as before the move.

#### Swap event ends between the tree vertices

Let *e*(*l*_0_, *l*_1_) denote unique event created from loci *l*_0_, *l*_1_, i.e., *e*(*l*_0_, *l*_1_) is equal to (*l*_0_, *l*_1_) if *l*_0_ *< l*_1_ and to (*l*_1_, *l*_0_) otherwise.

We sample two vertices – (*l*_0_, *l*_1_), (*u*_0_, *u*_1_) from *V*_*T*_ uniformly. If the vertices represent events that belong to different chromosomes, the move is rejected. Otherwise, we have four possible ways of obtaining new events for vertices (*l*_0_, *l*_1_), (*u*_0_, *u*_1_):

- *e*(*l*_0_, *u*_0_), *e*(*l*_1_, *u*_1_),
- *e*(*l*_0_, *u*_1_), *e*(*l*_1_, *u*_0_),
- *e*(*l*_1_, *u*_1_), *e*(*l*_0_, *u*_0_),
- *e*(*l*_1_, *u*_0_), *e*(*l*_0_, *u*_1_).

We sample one of those uniformly. If both new events are valid and belong to *V*_0_ then we change (*l*_0_, *l*_1_) and (*u*_0_, *u*_1_) to new labels with probability given by (13). Otherwise, the move is rejected.

Formally, if one of the new events is either not valid or is already present on the tree then we set the likelihood *P* (*T* ′, *θ*|*D, C*) of such structure to zero. This allows us to reject such a proposal deterministically since the acceptance ratio (13) is equal to zero.

This move changes both the *V*_*T*_ and *V*_0_ sets, but their cardinality remains the same. The move is symmetric.

#### Add leaf

We uniformly sample a vertex *v* from the set of inactive events *V*_0_ and add it as a leaf to uniformly sampled vertex from the *V*_*T*_ set. Denote the updated set of leaves by *V*_*l*′_. |*V*_*l*′_ | depends on the fact whether we added *v* as a leaf under internal vertex of the initial tree *T*, thus increasing the number of leaves, or under a leaf, leaving the number of leaves unchanged. To reverse this move, we resample *v* from *V*_*l*′_, remove *v* from the tree and add it to *V*_0_. This yields the Hastings ratio

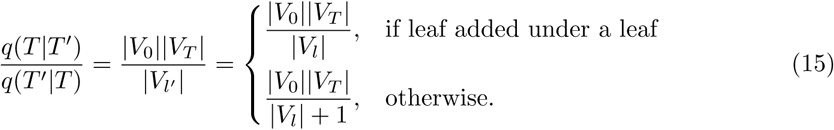

#### Remove leaf

We uniformly sample a leaf *v* from the set of leaves *V*_*l*_, remove *v* from the tree and add it to the set of inactive events *V*_0_. To reverse this move, we resample *v* from updated set of inactive events *V*_0′_ (*V*_0′_ = |*V*_0_| ∪ *v*), and add it as child to it’s former parent. To choose this parent, we sample it from the updated *V*_*T*′_ = *V*_*T*_ *\ v*. Consequently, the Hastings ratio for this move becomes

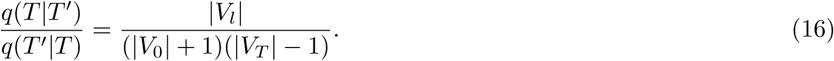

### Moves on parameter space

Recall that the vector of parameters is equal to:

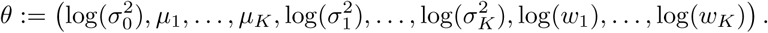

We use the Metropolis-within-Gibbs algorithm, i.e. one move from *θ* to *θ*′ consists of changing the value of only one of the coordinates. We use the deterministic scan strategy. The coordinate, which will be updated is chosen periodically – *i*-th step proposes new value for 1 + (*i* mod 3*K* + 1) coordinate. For every coordinate, we use adaptive scaling random walk Metropolis kernel, where the step size is adjusted to achieve optimal acceptance probability. [57].

### Improving the convergence of MCMC sampling

To improve the convergence of our algorithm we divide the sampling scheme into two chains. We start with a chain that works on the joint space (*T, θ*) and outputs the estimated maximum *a posteriori* value of mixture parameters (let us denote this value by *θ*\*). This chain alternates between moves changing *T* and moves changing *θ*, as was described in section MCMC Sampling. The second chain runs a fixed number *R* (this constant is user defined) of copies of the chain on the space of trees in parallel. The values of the mixture parameters are fixed and set to *θ*\*.

Let us describe the latter phase more precisely. Denote trees of the chains by *T*_1_, *…, T*_*R*_ and let *γ*_1_ = 1 > *γ*_2_ > *…* > *γ*_*R*_ > 0 be a sequence of *temperatures*. Chain number *i* targets distribution given by log-density

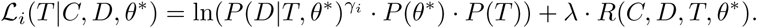

Notice that the difference between the standard distribution 12 and *ℒ*_*i*_ is that the likelihoods *P* (*D*|*T, θ*) are tempered and values of the parameters are held fixed.

One iteration of the second phase consists of two procedures:

- Each chain samples new tree 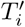 independently using scheme from MCMC Sampling,
- States of chains *J, J* + 1 are swapped with probability 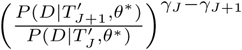. *J* is a random index sampled uniformly from {1, *…, R −* 1}.

This scheme is an example of *parallel tempering* algorithm [58] and it can be shown that its stationary distribution is proportional to 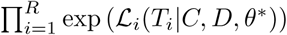. Instead of fixing specific values for the temperatures, we utilize the adaptive Parallel Tempering algorithm of [58]. Maximum *a posteriori* tree found by the latter scheme together with *θ*\* is defined to be the output of the whole procedure.

The speed of convergence of chain inferring *θ*\* is highly dependent upon a good choice of initial values for *θ*. We aggregate *D* matrix into one vector and assume that it is a sample from a mixture of *K* + 1 truncated normal distributions with non-negative means from which one is constrained to have mean zero. We employ the EM algorithm [59] to infer the parameters of this mixture and use them as initial values for *θ*.

## Simulated data

Simulations are performed to test of our model’s performance in the conditions where we know the ground truth CONET tree. Different simulation settings are generated, by varying the size *t* of the simulated tree *T*, the number of loci |*L*|, the number of single cells *m* and whether the distribution of the corrected count absolute differences for the breakpoint loci is well separated from the distribution for the non-breakpoint loci (*well separated setting*) or not (*poorly separated setting*). The data is simulated as originating from one chromosome.

First, the tree structure *T* is sampled uniformly from the set of all trees of pre-specified size *t*. The size *t* is either 20 with the number of all possible breakpoints |*L*| equal to 40 or two times more, i.e., 40 with 80 possible breakpoints. Next, we sample the events corresponding to each vertex from the set of all possible events with log-probability proportional to the negative length of the event. The probability of a cell being attached to a given vertex *v* is equal to 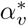, where 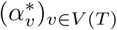 is sampled from Dirichlet(𝟙).

Finally, we generate simulated corrected count absolute differences *d*_*j,i*_ for each cell *j* = 1, *…, m* and each locus *i* ∈ *L*. To this end, for each cell, its breakpoints are read from its tree attachment. Differences for loci without breakpoint are sampled from distribution *f*_0_ while differences for loci with breakpoint are sampled from distribution *f*_*bp*_. In the *well separated setting* case we set:

- *f*_0_ = 𝒩 (0, 0.3)
- *f*_*bp*_ is a mixture of (𝒩 (1, 0.4), 𝒩 (2, 0.4), 𝒩 (3, 1.7)) with weights (0.5, 0.35, 0.15). In the *poorly separated setting* case we set:
- *f*_0_ = 𝒩 (0, 0.7)
- *f*_*bp*_ is a mixture of (𝒩 (1, 0.7), 𝒩 (2, 0.7), 𝒩 (3, 1.7)) with weights (0.5, 0.35, 0.15). Where 𝒩 (*a, b*) denotes normal distribution with mean *a* and standard deviation *b*. In the simulations we do not produce corrected count data *C* and thus set the *s*_0_ constant to 0, i.e., the *R*(*C, D, T, θ*) penalty is ignored in the inference.

### Model evaluation methods for simulated data

To facilitate the assessment of the quality of inference results for simulated data we introduce six performance scores. The scores can be divided into two groups – those that evaluate the quality of breakpoint detection and those that evaluate the quality of the inferred event history.

The breakpoint detection scores are based on the comparison of the inferred break-point matrix to the real (simulated) one. By breakpoint matrix we mean a matrix

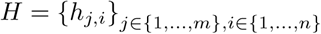

where *h*_*j,i*_ is equal to 1 when locus *i* is a breakpoint for cell *j* and 0 otherwise.

*False positive rate (FPR)* – number of 1 entries from inferred breakpoint matrix which are equal to 0 in the real matrix divided by the total sum of entries of the inferred matrix. FPR ranges from 0 to 1.

*False negative rate (FNR)* – number of 0 entries from inferred breakpoint matrix which are equal to 1 in the real matrix divided by the total sum of entries of the real matrix. FNR ranges from 0 to 1.

*Symmetric distance (SD)* – *L*_1_ distance between real and inferred breakpoint matrices divided by the number of cells. Notice that although the first two scores are divided by the total number of breakpoints, this one is not. As a result values of SD may exceed 1.0. For instance, *SD* = 1 indicates that on average one breakpoint is missed or wrongly inferred for each cell.

Lower values of *FPR, FNR* and *SD* indicate better breakpoint detection.

The event history scores facilitate the comparison of the inferred tree structure to the real (simulated) structure and consist of:

*Tree size (TS)* – the size of the inferred tree |*T* |, i.e. the number of vertices including root. In the evaluation, this number is compared to the real tree size.

*Edge sensitivity (ES)* – the proportion of the inferred tree edges that are present in the real tree.

*Edge precision (EP)* – the proportion of real tree edges that are present in the inferred tree.

The values of both *ES* and *EP* range between 0 and 1, where larger values indicate better inference of the event history.

### Model evaluation methods for biological data

Since in the case of biological data the ground truth about CONET structure is unknown, we introduce additional methods to assess the quality of our inference procedure applied to experimental data sets.

For basic characterization of the inferred trees we report *Tree size* as defined in Model evaluation methods for simulated data and *No of clusters* – the number of all *BC*_*w*_ clusters i.e. subsets of bins that share the same copy number event history, including the *B*_ø_ cluster.

To evaluate the consistency of an inferred CONET with the count data, we use characteristics defined below. Cluster support refers to a cluster’s inferred CN support calculated as the number of bins in a given cluster with corrected count close to inferred CN (*c* ∈ [*CN −* 0.5, *CN* + 0.5]) divided by the number of all the bins in the cluster.

*Avg cluster support* – average over all *BC*_*w*_ clusters’ supports,

*Perc of good clusters* – percentage of clusters with at least 0.7 support,

*Perc of CN=2 clusters* – percentage of clusters that infer CN equal to two (not including *B*_ø_ cluster).

Finally, we define detailed measures evaluating the quality and consistency of the CN calling results, where by bins inside events we mean bins in all *BC*_*w*_ clusters except for *BC*_ø_.

*Avg Gini Index in events* – Gini Index calculated separately for corrected counts in bins for each *BC*_*w*_ cluster (*w* ≠ ø), averaged over the clusters,

*Avg Entropy in events* – normalized Shannon Entropy calculated separately for corrected counts in bins in each *BC*_*w*_ cluster (*w* ≠ ø), averaged over the clusters; to calculate the entropy in each cluster the corrected counts are grouped into intervals of width 1, concentrated around integer copy numbers,

*Gini Index outside events* – Gini Index calculated for corrected counts in bins in the *BC*_ø_ cluster,

*Entropy outside events* – normalized Shannon Entropy calculated for corrected counts in bins in the *BC*_ø_ cluster,

*CN-TPR* – The number of bins inside events with corrected count *c* far from two, i.e. *c* ∉ [1.5, 2.5], divided by the total number of bins with the true corrected count far from two,

*CN-FPR* – The number of bins inside events with corrected count *c* close to two, i.e. *c* ∈ [1.5, 2.5]m divided by the total number of bins with the true corrected count close to two.

*CN-RMSE* – root mean square error for CN calling, the quadratic mean of the differences between the inferred integer copy number for each bin and the corrected count for each corresponding bin, which is equal to the square root of count discrepancy penalty: 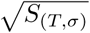

## Availability of data and materials

CONET implementation is freely available under CC-BY-NC 4.0 International license at https://github.com/tc360950/CONET.

## Ethics declarations

### Ethics approval and consent to participate

Not applicable.

### Consent for publication

Not applicable.

### Competing interests

Merck Healthcare KGaA provides funding for the research group of ESz. MM is co-financed by part of these research funds.

## Author’s contributions

MM and ESz conceptualized the initial idea of the model. MM, TC, ESz and BM developed the methodology. MM implemented the draft version of the model for cross-checking, TC implemented and optimized the final version. TC prepared the simulated data and validation, MM processed the biological data and performed biological validation. MM and TC prepared the Figures. MM, TC and ESz wrote the initial draft of the manuscript. All authors provided critical feedback; helped shape the research and analysis; edited, reviewed and approved the manuscript.

## Acknowledgements

The work by MM was done during Interdisciplinary doctorate studies using next generation sequencing in personalized medicine (Interdyscyplinarne Studia Doktoranckie wykorzystujące sekwencjonowanie nowej generacji (NGS) w medycynie spersonalizowanej). MM is grateful for discussions during her secondment at Dr. Shah’s group at the Memorial Sloan Kettering Cancer Center.

## Funding

The work was funded by the Polish National Science Centre grant OPUS no. 2019/33/B/NZ2/00956 to ESz, and by the Polish National Science Centre grant OPUS no. 2018/31/B/ST1/00253 to BM. MM was co-funded by the European Social Fund POWER programme (financing agreement no. POWR.03.02.00-00-I041/16-00).

## Additional Files

Additional file 1

List of the inferred CONET vertices for xenograft SA501X3F breast cancer sample.

## Notes

https://github.com/tc360950/CONET

### Competing Interest Statement

Merck Healthcare KGaA provides funding for the research group of Ewa Szczurek. Magda Markowska is co-financed by part of these research funds.

